# Pathogens and Antimicrobial Resistance Genes in Household Environments: A Study of Soil Floors and Cow Dung in Rural Bangladesh

**DOI:** 10.1101/2024.12.06.627269

**Authors:** Anna T. Nguyen, Kalani Ratnasiri, Gabriella Barratt Heitmann, Sumaiya Tazin, Claire Anderson, Suhi Hanif, Afsana Yeamin, Abul Kasham Shoab, Ireen Sultana Shanta, Farjana Jahan, Md. Sakib Hossain, Zahid Hayat Mahmud, Mohammad Jubair, Mustafizur Rahman, Mahbubur Rahman, Ayse Ercumen, Jade Benjamin-Chung

**Affiliations:** Department of Epidemiology & Population Health, Stanford University; Stanford Immunology Program, Stanford University School of Medicine, Stanford, CA 94305, USA; Department of Forestry and Environmental Resources, North Carolina State University; Department of Civil & Environmental Engineering, Stanford University; Environmental Health and WASH, International Centre for Diarrhoeal Disease Research, Bangladesh; Emerging Infections, Infectious Diseases Division, International Centre for Diarrhoeal Disease Research, Bangladesh; Laboratory of Environmental Health, International Centre for Diarrhoeal Disease Research, Bangladesh; Genomics Centre, International Centre for Diarrhoeal Disease Research, Bangladesh; Global Health and Migration Unit, Department of Women’s and Children’s Health, Uppsala University, Sweden; Chan Zuckerberg Biohub San Francisco

## Abstract

In low- and middle-income countries, living in homes with soil floors and animal cohabitation may expose children to fecal organisms, increasing risk of enteric and antimicrobial-resistant infections. Our objective was to understand whether cow cohabitation in homes with soil floors in rural Bangladesh contributed to the presence and diversity of potential pathogens and antimicrobial resistance genes (ARGs) in the home. In 10 randomly selected households in rural Sirajganj District, we sampled floor soil and cow dung, which is commonly used as sealant in soil floors. We extracted DNA and performed shotgun metagenomic sequencing to explore potential pathogens and ARGs in each sample type. We detected 6 potential pathogens in soil only, 49 pathogens in cow dung only, and 167 pathogens in both soil and cow dung. Pathogen species with relative abundances >5% in both soil floors and cow dung from the same households included *E. coli* (N=8 households), *Salmonella enterica* (N=6), *Klebsiella pneumoniae* (N=2), and *Pseudomonas aeruginosa* (N=1). Cow dung exhibited modestly higher pathogen genus richness compared to soil floors (Wilcoxon signed-rank test p=0.002). Using Bray-Curtis dissimilarity, pathogen species community composition differed between floors and cow dung (PERMANOVA p<0.001). All soil floors and cow dung samples contained ARGs against antibiotic classes including sulfonamides, rifamycin, aminoglycosides, lincosamides, and tetracycline. Paired floor and cow dung samples shared ARGs against rifamycin. Our findings support the development of interventions to reduce soil and animal feces exposure in rural, low-income settings.

**Importance:** In low-income countries, inadequate housing materials and animal cohabitation can lead to fecal contamination of rural homes. Contaminated soil floors are difficult to clean and may harbor organisms causing illness and antibiotic resistance, especially in young children, who frequently ingest soil. We sequenced soil floor and cow dung samples from households in Sirajganj district, Bangladesh and identified pathogens and antibiotic resistance genes. We detected 167 pathogens in both soil and cow dung; pathogens present in both sample types at the highest relative abundances were *E. coli*, *Salmonella enterica*, *Klebsiella pneumoniae,* and *Pseudomonas aeruginosa*. Antibiotic resistance genes were found in all samples. In cow dung, the most common genes conferred resistance to the antibiotics lincosamide, rifamycin, cephamycin, and tetracycline. In soil floors, the most common genes conferred resistance to rifamycin, sulfonamides, and aminoglycosides. Household soil and cow dung may be important reservoirs of pathogens and antimicrobial resistance in low-income countries.

## Introduction

In low- and middle-income countries (LMICs), inadequate housing remains common^1,2^ and is associated with disease and mortality^3^. Rural households frequently cohabitate with domestic animals in LMICs, and animal husbandry is a critical source of income and nutrition^4^. Yet, animals also contribute to fecal contamination of rural households^5,6^. Studies using avian and ruminant microbial source tracking markers found that animals contribute to *Escherichia coli* – a widely used environmental indicator of human fecal contamination – in soil in households in Bangladesh^7^. Young children frequently touch and ingest soil, which may be contaminated with human or animal feces, within household premises^8–10^. A prior study in Bangladesh estimated that mouthing of child hands, direct soil ingestion, and direct feces ingestion were leading contributors to child *E. coli* ingestion among children under 1 year in household settings^9^.

Young children’s exposure to soil and animal feces can facilitate disease transmission in the home. Household soil floors are a largely overlooked reservoir for soil-transmitted helminths (STH), *Shigella*, pathogenic *E. coli* ^11–13^, and possibly other pathogens. Some studies have detected levels of *E. coli* and STH in household soil floors that exceed those in samples taken from latrine floors^12,14,15^. Contact between young children and domestic poultry and livestock is associated with an increased risk of diarrhea^16^. Exposure to household fecal contamination via household soil, stored drinking water, child hands, and fomites, is associated with increased risk of diarrhea^17^, enteric pathogen infections^18^, child growth faltering^17,18^, and antimicrobial resistance^19^.

Soil harbors diverse bacteria with naturally occurring antibiotic resistance that can exchange genes or plasmids with human and animal pathogens in fecally contaminated soils, making such soils an important reservoir of emerging antimicrobial resistance^20–22^. While it has been suggested that ARGs likely are transmitted between soil and human pathogens^20^, studies have yet to demonstrate this^23^. Prior studies have detected extended-spectrum beta-lactamase (ESBL)-producing *E. coli* in household soil in rural Bangladesh^24^ and multidrug resistant *E. coli* in household yard soil in Tanzania^25^. A genomic study found that multiple *E. coli* genes in rural Bangladesh were associated with virulence and antibiotic resistance in household and yard soil, and phylogenetic analyses suggested that *E. coli* in soil was likely from diverse human and animal sources^19^.

Studies have also found evidence of horizontal gene transfer of antimicrobial resistance genes between animals and humans in household settings in LMICs^26^. Unhygienic animal husbandry practices (e.g., handling cow dung with bare hands), sharing of household spaces with domestic animals, poor management of animal waste and carcasses, and inadequate hygiene while caring for domestic animals contribute to zoonotic and antimicrobial-resistant (AMR) pathogen transmission in LMICs^27,28^. Moreover, inadequate access to veterinary resources contributes to the misuse of antimicrobials for prophylaxis or as feed additives in low income settings^29^. A recent review of studies investigating the contribution of animals to AMR in humans reported mixed results ^30^; they concluded that this question remains poorly understood in LMICs because many prior studies were either conducted in high-income settings with low levels of human-animal contact or did not use appropriate methods to capture transmission dynamics.

In rural, low-income communities where cattle rearing is common, cows often cohabitate closely with humans, and cow dung may contribute to household pathogen contamination. Cow dung is commonly used as fertilizer, cooking fuel, and as a coating for floors and household walls in household settings in LMICs and in Bangladesh^27,31–33^, yet the extent to which household contamination with cow dung contributes to zoonotic or AMR pathogen transmission is unknown. Cow dung commonly contains human pathogens such as *Salmonella spp.*, *Campylobacter spp.*, *Listeria monocytogenes*, *Yersinia enterocolitica*, *E. coli*, *Cryptosporidium parvum* and *Giardia lamblia*^34^. A prior study in Bangladesh found that the presence of cow dung in household courtyards and detection of a molecular marker of cow feces on mothers’ hands was associated with the presence of pathogenic *E. coli* and *Giardia* on mothers’ hands^6^. Carbapenem-resistance genes^35^ and AMR *E. coli* and *Salmonella spp.* – WHO high-priority pathogens for development of AMR^36^ – have been detected in cow dung samples^37^. Studies in Bangladesh have also found high levels of carbapenem-resistance genes in household cattle dung samples^35^.

Our objective was to understand whether cow cohabitation in homes with soil floors in rural Bangladesh contributed to pathogen and antimicrobial resistance genes (ARGs) in the household setting. We hypothesized that 1) cow dung and soil floors would contain human pathogens and ARGs and 2) cow dung and soil floors from the same households would have overlapping microbiomes and resistomes. We conducted this exploratory pilot study in rural Sirajganj district, Bangladesh, where cattle rearing is common, and cows frequently cohabitate inside homes with soil floors.

## Results

We enrolled 10 households from Sthal union, Chauhali upazila, Sirajganj Bangladesh. Households were eligible for enrollment if they had a soil floor, a child under the age of 2 years, available cow dung for sampling and no self-reported cases of anthrax among their domestic animals or household members. The mean number of household members was 6 (range 4-8), and most homes were approximately 300 square feet (Table 1). Households typically had access to a pit latrine and tubewell within their compound. In addition to keeping cows, all but one household owned sheep or goats, and all but one owned chickens, ducks, or pigeons. The mean number of animals owned by each household was 4 cattle, 4 sheep or goats, and 17 chickens, ducks, or pigeons. Seven households kept their cows tied up outside during the day, while the rest kept them in a different structure in the compound (e.g., a cattle shed). At night, seven households kept their cows inside a household while the rest kept them in a different structure in the compound or tied up outside (Table A1). Four of ten households use cow dung for household purposes. At the time of the survey, wet, dry, or processed cow dung was visible in the courtyard of eight households and on the main area of indoor household floors in two households.

**Table 1.** Characteristics of study households

|  | Mean (Range) or N (%) |
| --- | --- |
| Number of household members | 6.3 (4, 8) |
| Number of children under 5 in the household | 1.3 (1, 3) |
| Number of animals owned |  |
| Cattle | 3.6 (1, 8) |
| Goat or sheep | 3.5 (0, 7) |
| Chickens, ducks, or pigeon | 16.8 (0, 30) |
| Location of cattle during the daytime |  |
| In a different house in the compound | 3 (30%) |
| Tied up outside | 7 (70%) |
| Location of cattle at nighttime |  |
| Free inside the home with no barrier | 2 (20%) |
| Tied up inside the home with barrier | 5 (50%) |
| In a different house in the compound | 1 (10%) |
| Tied up outside | 2 (20%) |
| What is done after cow defecates |  |
| Remove it from the house | 1 (10%) |
| Remove it from the house and then clean the floor | 7 (70%) |
| Pile it inside the house | 2 (20%) |
| Household uses cow dung for cooking, fertilizer, etc. | 4 (40%) |
| Cow dung visible in the courtyard | 8 (80%) |
| Cow dung visible on the main area of the household floor | 2 (20%) |

### Microbes

We performed shotgun metagenomic sequencing and used Kraken/Bracken for taxonomical analyses on paired floor soil and cow dung samples from the same households. Sequencing yielded a total of 152.94 million reads of DNA from the 10 cow dung samples (7.18-28.4 million reads per sample) and 33.28 million reads from the 10 soil samples (0.58-11.01 million reads per sample) (Table A2). After quality filtering for all samples and host filtering for cow dung samples, the average reads remaining per sample were 3.6 million in cow dung (0.1-10.4 million reads per sample; 22.6% retained on average) and 1.2 million in soil (0.3-3.5 million reads per sample; 39.9% retained on average) that were used as the input for metagenomic analysis.

Soil and cow dung samples exhibited a large number of microbes, with an average of 1,935 genera and 6,799 species detected per sample in cow dung and 1,057 genera and 3,837 species detected per sample in soil floors. The genera with the highest relative abundance were *Bacteroides*, *Faecalibacterium*, *Prevotella, Clostridium*, and *Phocaeicola* in cow dung and *Janibacter, Nocardioides, Streptomyces, Brachybacterium,* and *Serinicoccus* in soil.

In order to focus on microbes with pathogenic potential, we restricted microbial analyses to microbes with known pathogenicity in humans^38^. Across potential pathogens, the most abundant genera across samples were *Prevotella*, *Clostridium*, *Bacillus*, *Pseudomonas*, and *Streptococcus* in cow dung samples and *Pseudomonas*, *Mycolicibacterium*, *Corynebacterium*, *Escherichia*, and *Mycobacterium* in soil floors. The most common pathogen species in cow dung were *Clostridioides difficile*, *E. coli* (including pathogenic and/or non-pathogenic strains), *Salmonella enterica*, *Klebsiella pneumoniae*, and *Prevotella melaninogenica;* in soil, the most common pathogens species were *E. coli*, *K. pneumoniae*, *S. enterica*, *Pseudomonas aeruginosa*, and *Pseudomonas putida* (Figure 1). We detected 6 potential pathogens in soil only, 49 pathogens in cow dung only, and 167 pathogens in both soil and cow dung. Pathogen species with relative abundances >5% in both soil floors and cow dung from the same households included *E. coli* (N=8 households), *S. enterica* (N=6), *K. pneumoniae* (N=2), and *P. aeruginosa* (N=1) (Figure 2, Figure A1).

**Figure 1.**
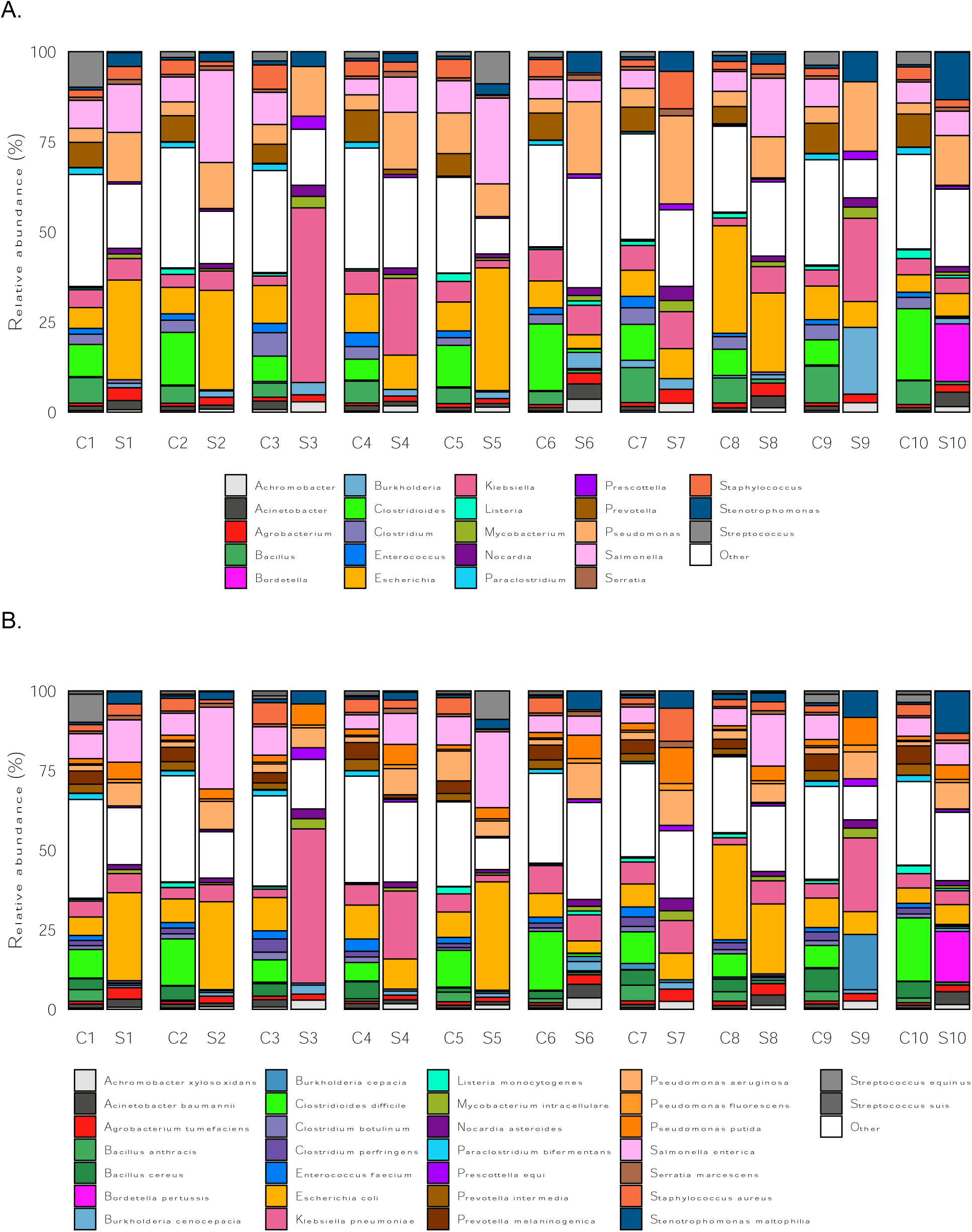
Sample-level relative abundance of non-host reads for potential pathogens by mNGS analysis at the A) genus-level and B) species-level. Includes the top 30 species by average relative abundance across all samples, with all other genera or species labeled as “Other.” C1-C10 refer to cow dung samples, and S1-S10 refer to floor soil samples; each number corresponds to a different household (e.g., C1 and S1 are from household 1).

**Figure 2.**
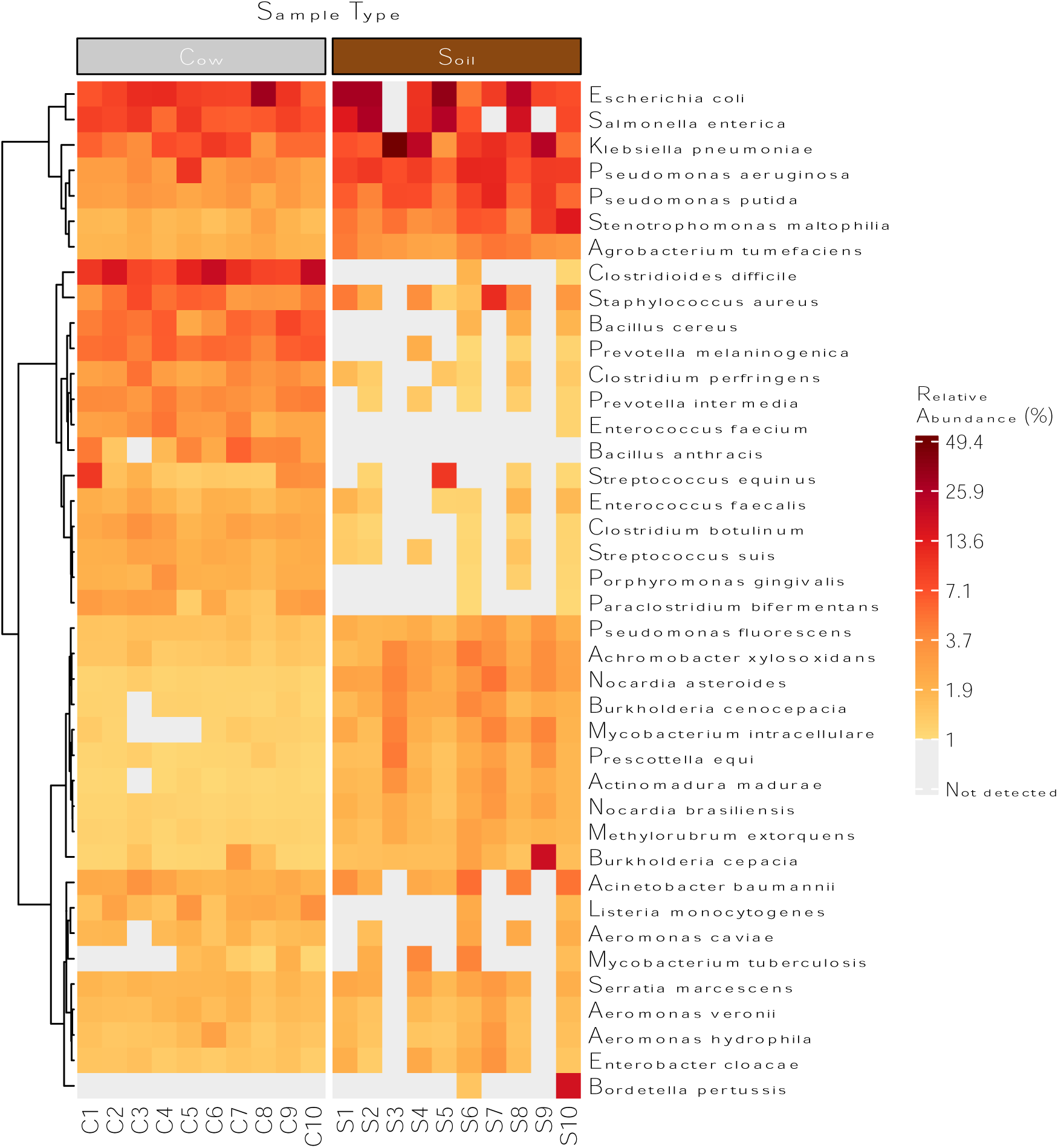
Heatmap of filtered, non-host reads for potential pathogens detected in samples of cow dung (listed as C1-C10) and floor soil (listed as S1-S10). Tile colors indicate the relative abundance of each species within each sample. Gray tiles indicate that a species was not detected. Includes taxa with an average relative abundance across all samples of at least 0.5%. The heatmap displays hierarchical clustering of rows using Euclidean distance and Ward’s minimum variance method.

### Diversity of potential pathogens within samples

We determined the alpha-diversity of potential pathogens using measures of microbial genus richness and evenness within each sample type using: the richness attribute, Chao1 index for species richness^39–41^, Pielou’s evenness index^42^, Shannon index^43^, Simpson index^44^, and inverse Simpson index. Cow dung samples exhibited modestly higher levels of pathogen species and genus richness (Wilcoxon signed-rank test p=0.002 for species, p=0.002 for genera) compared to soil samples (Figure 3, Figure A1). Patterns were similar for pathogen species diversity by Chao1. There was a wider range of within-sample pathogen diversity in soil samples compared to cow dung samples by diversity metrics that included evenness.

**Figure 3.**
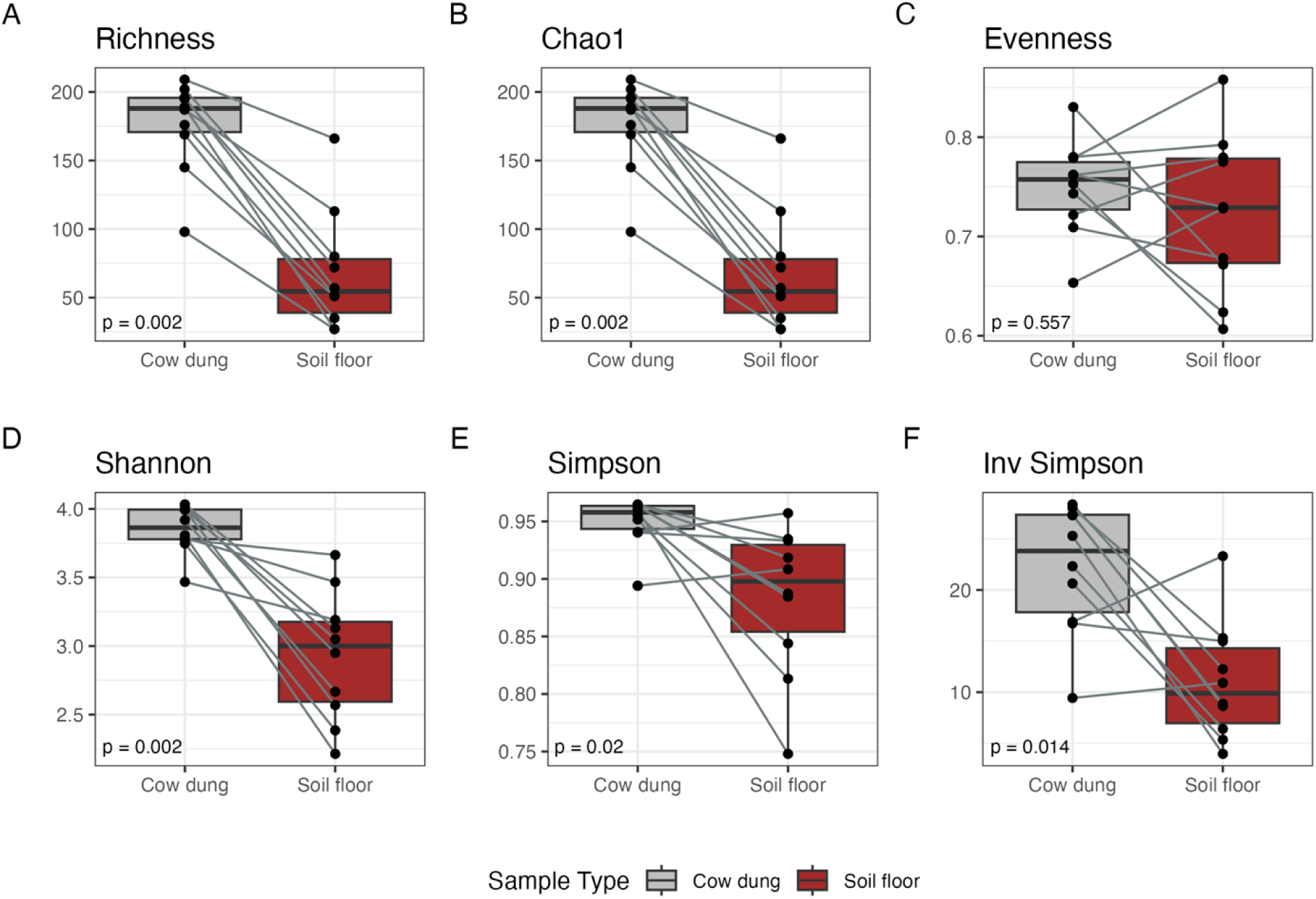
Alpha-diversity indices for potential pathogen species by sample type. Includes 10 household-paired cow dung and soil floor samples. Indices were compared between sample types using the Wilcoxon signed-rank test.

### Diversity of potential pathogens between samples

To compare diversity of potential pathogens between samples, we calculated Bray Curtis dissimilarity and performed principal coordinates analysis. Potential pathogen species community composition differed between floor and cow dung samples (pairwise permutational multivariate analysis of variance R^2^=0.54, p<0.001) but not between households (R^2^=0.22, p=0.998) or between households with and without visible animal feces on the floor inside the home (R^2^=0.01, p=0.9498) (Figure A2).

### Antimicrobial resistance

Next, we detected antimicrobial resistance genes (ARGs) using the Chan Zuckerberg ID (CZID) pipeline. Analyses revealed diverse ARG profiles with substantial variation between households and sample types. The most common ARGs we detected conferred resistance to rifamycin, sulfonamide, aminoglycoside, or multiple drug classes in soil floors and tetracycline, cephamycin, lincosamide, rifamycin, or multiple drug classes in cow dung (Figure 4, Figure 5a). ARGs against beta-lactam, lincosamide, tetracycline, cephamycin, rifamycin, and multiple drug classes were present in at least half of cow dung samples; ARGs against mupirocin-like antibiotic, sulfonamide, aminoglycoside, and multiple drug classes were present in at least half of soil floor samples. Only ARGs that confer resistance to rifamycin were found in both soil floors and cow dung in multiple households. There was a larger number of distinct ARGs in soil floors than in cow dung samples. Genes associated with resistance to multiple classes of antibiotics were common in both sample types, particularly in soil. ARGs in soil most commonly conferred resistance through antibiotic target protection, antibiotic inactivation, and target alteration or replacement; the most common mechanism for ARGs in cow dung was antibiotic efflux (Figure 5b). Nine of 10 soil floors and 9 of 10 cow dung samples contained at least one ARG in the highest quartile of risk to human health (e.g., sul1, tet(Q), mexF, ermF, cfxA2) (Figure A3) (42).

**Figure 4.**
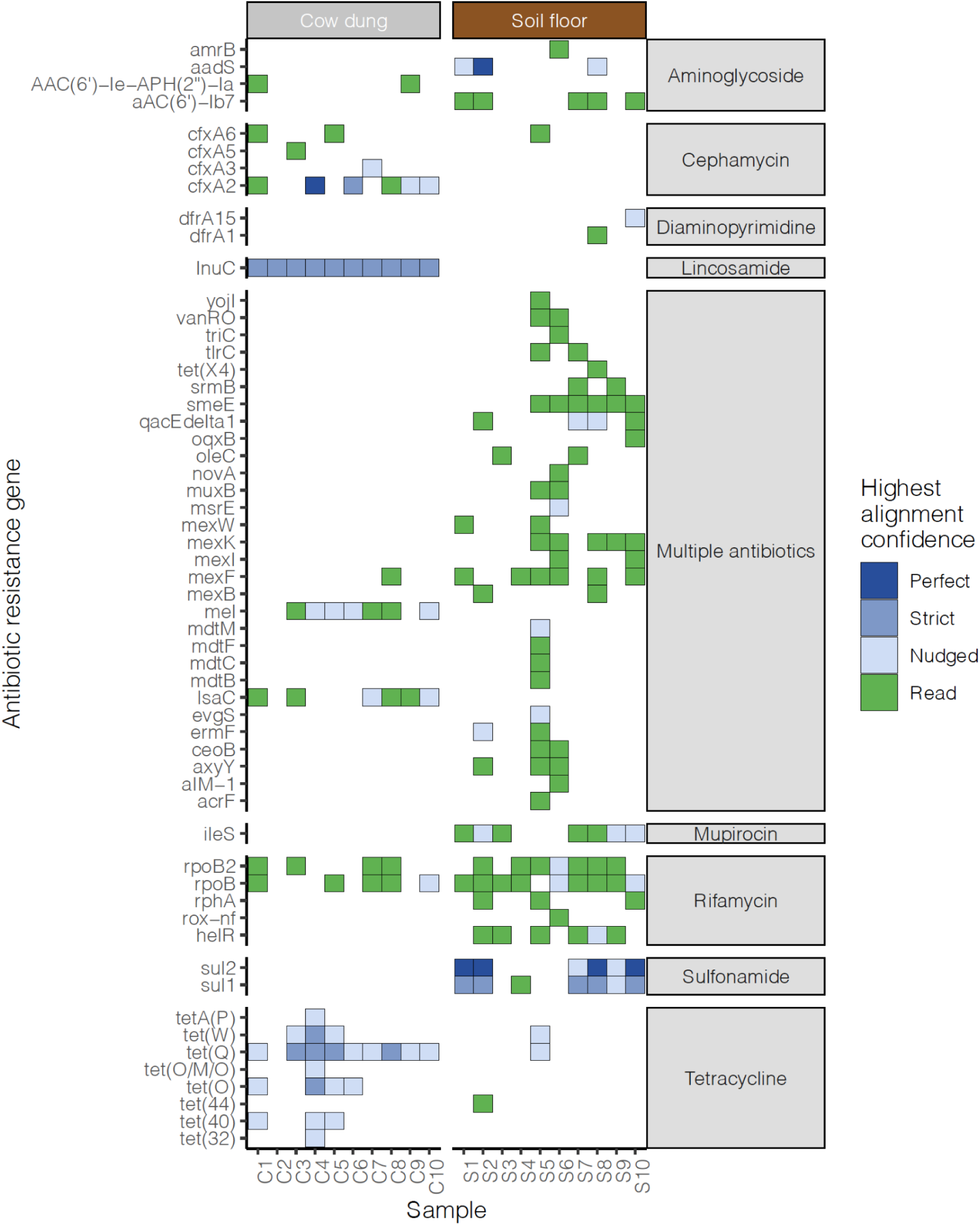
Heatmap of antibiotic resistance genes detected in cow dung and soil samples. Tile colors indicate the read coverage breadth. Includes genes with read coverage breadth > 10% or contig coverage breadth > 10% and > 5 reads mapped. Right annotation indicates the drug class that the ARG confers resistance to. Colors indicate the highest alignment confidence based on contig match quality (blue) or reads (green). “Perfect” contig matches identically matched reference sequences in the Comprehensive Antibiotic Resistance Database. “Strict” contig matches were those that matched previously unknown variants of known ARGs, including secondary screening for key mutations. “Nudged” contig matches had at least 95% identity to known AMR genes and were matched using a percent identity threshold not taking alignment length into account.

**Figure 5.**
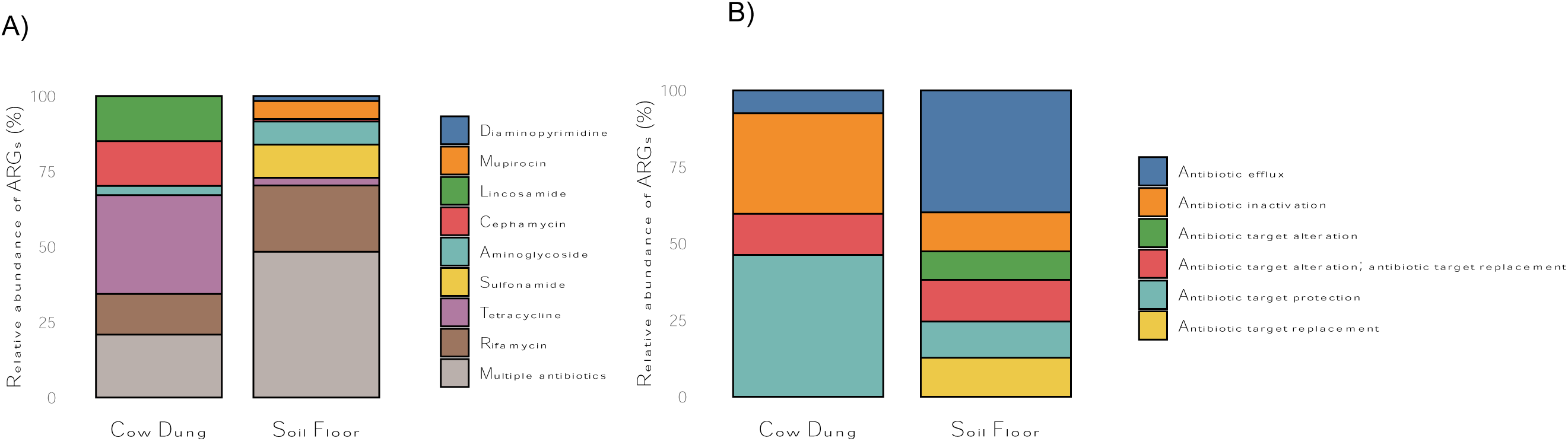
Relative abundance of antibiotic resistance genes detected in cow dung and soil samples by A) drug class and B) antibiotic resistance mechanism. Both panels include genes with read coverage breadth > 10% or contig coverage breadth > 10% and > 5 reads mapped.

## Discussion

In this exploratory study, we found that household soil floors and cow dung in rural Bangladesh contained diverse microbial communities, including numerous potential human pathogens. While our analysis does not allow us to infer transmission from cow dung to soil floors, our findings suggest that the presence of cow feces in domestic spaces may contribute to microbial contamination of household surfaces. Antimicrobial resistance genes against multiple drug classes were prevalent in both sample types, and nearly all soil floor and cow dung samples contained ARGs associated with increased risk to human health^45^. There were few shared ARGs present in paired samples from the same households. However, our small sample size may have limited our ability to establish a link between these sample types. Given children’s high levels of soil and animal contact in this setting^10^, our findings suggest that household soil floors and cow dung may be important reservoirs of diverse pathogens and antimicrobial resistance.

*E. coli* and *S. enterica* were present in both sample types in higher relative abundances, suggesting that soil and cow dung may be a household source of enteric infections. Many *E. coli* strains are commensal, and while our limited sequencing depth did not allow for pathogenic *E. coli* strain identification, a separate analysis of this study population found that 8% of *E. coli* isolates were pathogenic.^46^ Other studies in Bangladesh have detected a similar prevalence of pathogenic *E. coli* in household soil samples.^24,19^ While we are not aware of any prior studies in LMICs that have detected *Salmonella* in household floors, studies in the U.S. have identified *Salmonella* in soil near household entrances and in vacuum cleaners in homes with infected infants.^47,48^ In cattle, *S. enterica* is a facultative pathogen that can cause illness (e.g., enteric infection, reproductive loss); additionally, certain serotypes, such as *S. enterica* serotype Dublin, can result in lifelong asymptomatic carriage with intermittent shedding in cattle stool and severe illness if transmitted to humans.^49,50^

We also detected potential pathogens in soil floors and cow dung that can cause illness in individuals with lower immunity (*K. pneumoniae, P. aeruginosa, Stenotrophomonas maltophilia*, *C. difficile, Staphylococcus aureus*). The presence of pathogens in household environments has been documented in LMICs and high-income countries, though studies in LMICs are limited. A study in Malawi detected extended spectrum beta lactamase *K. pneumoniae* at low levels on household floors,^51^ while research in Ghana identified some of the same microbes we detected in household dust, including *Acinetobacter baumannii*, *Bacillus cereus*, and *Enterobacter cloacae*.^52^ *C. difficile* was present in all cow dung samples, consistent with other studies which have found that it can colonize healthy cattle and be transmitted zoonotically.^53^ It was only detected in two soil samples, consistent with a prior study that detected it in approximately one-third of household soil samples in rural Zimbabwe.^54^ In high-income countries, household surfaces can also be highly contaminated with potential pathogens; *E. coli, K. pneumoniae, P. aeruginosa, S. aureus,* and *S. maltophilia* have been found on sink and shower drains, floors, and surfaces.^55–58^ Additionally, studies in high-income countries has found that household dust contains *E. coli, Pseudomonas, Acinetobacter, Enterobacter, Enterococcus, Bacillus,* and *Staphylococcus*.^59^ The health implications of household contamination with these pathogens remain unclear, and it is not known whether the levels of contamination commonly seen in LMICs or high-income countries contribute to infections.^60^ Our metagenomic analysis did not allow us to quantify the concentration of potential pathogens; assessing the risk of infection under typical low-income country household exposures using culture-based methods and quantitative microbial risk assessment is an important area of future research.

Many potential pathogens were present in soil floors and cow dung samples from the same households, which may reflect contamination of floors with cow feces. However, our cross-sectional sample and use of metagenomic sequencing did not allow us to establish the source of microbes in soil floors. Prior studies in Bangladesh have detected ruminant fecal markers in household soil and hand rinses in households that both owned or did not own ruminants.^7,61^ Some of the pathogens found in soil floors are common in soil or other environmental niches exposed to human activity, so their presence may not imply fecal contamination by cow dung (e.g., *E. coli*, *K. pneumoniae*, *B. cereus*, *P. aeruginosa*^62–66^. Other pathogens are commonly found in the gastrointestinal tract of warm-blooded animals^67–69^, such as *E. coli, K. pneumoniae*, and *S. enterica*, so their presence in soil may suggest that floors were contaminated with feces of humans or other animals, such as chickens or goats. *B. cereus* and *P. aeruginosa* also cause mastitis in cattle^70,71^, and it is possible that they are shed in cow dung during infection. Future research using microbial source tracking and phylogenetic analyses with longitudinal samples could elucidate the contribution of cow feces to household soil microbiota.

There was limited overlap in ARGs in paired soil floor and cow dung samples, and ARGs in each sample type provided resistance to different drug classes. These findings may imply that cow fecal contamination did not result in transmission of ARGs to soil. However, there are also several other possible explanations: (1) transferred DNA may have degraded over time, (2) ARGs may have been present below detection limits, (3) soil bacteria may harbor their own distinct resistome shaped by local selective pressures, (4) soil ARGs may primarily originate from other sources (e.g., human or chicken feces), or (5) ARGs in cow dung may not reside on mobile genetic elements capable of transfer between bacteria.^72^ Additionally, soil is a rich reservoir of ARGs, deriving from both natural and anthropogenic processes. Some ARGs we detected in household soil have been detected in pristine soils in Tibet and thus may not reflect fecal pollution (e.g., vanRO, rpoB2, rpoB, rphA, muxB, mexF, mexK, mexW).^73^ Commensal bacteria in the cattle gut, such as *E. coli*, could also be reservoir of ARGs; prior studies have found high levels of AMR in *E. coli* isolates from cattle^74^ and have linked ARGs in *E. coli* in household soil in Bangladesh to human and animal sources^19^. Additionally, livestock manure has been identified as a primary source of ARGs in agricultural soils.^75^ Regardless of their source, the presence of ARGs in household environments could contribute to the spread of AMR because mobile genetic elements can transmit them from environmental reservoirs to human pathogens.^76^

Our analysis revealed ARGs in environmental samples that align with findings of prior studies on ARGs in environmental samples and local antibiotic usage patterns. The most common ARGs we detected in cow dung and soil confer resistance to tetracyclines, rifamycins, sulfonamides, lincosamides, and aminoglycosides. Evidence was strongest for ARGs against sulfonamides in soil samples and lincosamides and tetracyclines in cow dung. A systematic review of ARGs in agricultural soils identified ARGs against tetracyclines, sulfonamides, and aminoglycosides.^64^ ARGs resistant to rifamycin were relatively common in both sample types. Studies in dairy farms have detected *Salmonella* spp. and *Listeria* resistant to rifamycins.^77^ Additionally, many environmental bacteria are naturally resistant to rifamycin, such as *Amycolatopsis*, which was present in nearly all cow dung and soil samples. However, rifamycins have not been detected in prior studies of ARGs in environmental reservoirs.^78^ ARGs we detected conferred resistance to antibiotics that are commonly used in this population. In a survey in Mymensingh, Bangladesh, farmers commonly treated animals with tetracyclines, sulfonamides, and aminoglycosides^79^. Lincosamides and rifamycins are frequently used to treat mastitis in cows^77,80^. Tetracycline is commonly used to treat gram-positive and gram-negative bacteria in animals and humans and to promote growth in livestock^81^.

A strength of using metagenomic sequencing in environmental samples is that it can reveal more ARGs than culture-based approaches since many bacteria in environmental reservoirs cannot be cultured.^72^ However, culture-based approaches are required to assess functional resistance through bacterial growth inhibition. A separate analysis of this study population found cefotaxime-resistant *E. coli* in 71% of household floors, and all isolates produced extended-spectrum beta-lactamase (ESBL) and were multi-drug resistant^46^. The majority of isolates contained the bla_CTX-M_ gene, and some samples contained bla_TEM_ and bla_SHV_ genes. This analysis complements the prior study to show that household floors and cow dung contain a wide range of additional ARGs beyond those that inhibit beta lactams. Taken together, our findings underscore the potential contribution of household soil to emerging AMR in low-income settings with high levels of animal contact.

Our study also has some limitations. First, because metagenomic sequencing did not include control samples, there are limitations in determining background contamination potentially introduced during the sample collection and library preparation process that may lead to false positives. There are also limitations in the methodology of short-read shotgun metagenomic sequencing. It is difficult to determine the sensitivity of metagenomic Next-Generation Sequencing (mNGS) to pick up all present microbes and ARGs equally in a sample as well as clearly distinguish contaminants from sample-associated microbes. It is important to note that here we present the results of mNGS work as exploratory research and these results need to be further validated with techniques like PCR and culture-based methods. Second, our choice and use of the pathogen filter at the genus and species level may not be as inclusive of all known human pathogens or granular enough to decipher pathogenicity. For example, within a single microbe genera and species there may be some strains that are non-pathogenic while others are pathogenic (e.g., *E. coli*). Third, we only investigated pathogens in cows, but other studies have found that chickens are important for AMR in similar settings^82^. We also did not collect data on antibiotic use for cows, chickens and household members. Fourth, we only detected pathogens using DNA, so our study excluded RNA viruses. Because we used a cross-sectional design, we were not able to rigorously investigate the directionality of pathogen or ARG transmission between cows and soil. Additionally, as this was an exploratory pilot study, we did not include a control group (e.g., households with soil floors and no cows). Finally, our sample size was small, which limits the generalizability of our findings.

Our findings contribute to the growing literature on household soil and domestic animals as potentially important contributors to disease transmission and as reservoirs of antimicrobial resistance in low-income country settings. Overall, our finding that household soil floors harbored diverse pathogens and ARGs underscores the need for housing upgrades and animal management improvements in low-income settings. Future interventions to reduce infections and AMR in similar household settings may consider focusing on reducing exposure to soil and cow dung.

## Methods

### Sample collection

This study was conducted as pilot study as part of the CRADLE trial.^83^ The pilot study enrolled households in Sthal union of Chauhali sub-district in Sirajganj district, Bangladesh. Sthal is located in a rural area on a sand bar within the Jamuna River. It is highly susceptible to flooding and erosion. The community residing in Sthal is primarily composed of agricultural workers, and cattle rearing is common, and the majority of homes have soil floors. Field staff enrolled 10 households. Households were eligible for enrollment if they had a child under 2 years of age, a floor fully made of soil and not fully covered with a mat, carpet, or jute sack, and if cow dung was available for sampling. Because anthrax outbreaks have occurred in the area prior to the study, for the safety of the study staff, we further restricted to households with no cases of anthrax among domestic animals or household members. Cow dung and floor soil were collected from each household.

To collect floor soil, field staff placed a bleach-sterilized 50 cm x 50 cm metal stencil on the floor next to the head of the bed where the child under 2 years of age slept. Using a sterile scoop, they scraped the soil inside the stencil once vertically and then once horizontally, with the goal of collecting 20 g of soil. Soil was placed in sterile Whirlpak bags. Field staff collected fresh cow dung from a defecation event since dawn the same day. They prioritized collecting cow dung from the same room as the floor samples. If this was not possible, they collected it from another room in the house, and if that was not possible, they collected it from the compound courtyard or the edges of the compound. Cow dung was collected using a sterile stool collection tube with a sterile spoon. Floor scrapes and cow dung were placed in a cooler with ice and transported to the International Centre for Diarrhoeal Disease Research, Bangladesh (icddr,b) laboratory in Dhaka for analyses. All samples were stored at 4°C overnight and transferred to -80°C freezer the following morning to be stored until DNA extraction.

### DNA extraction

DNA was extracted from 250 mg aliquots of cow dung using the QIAamp PowerFecal Pro DNA Kit (50) (Cat# 51804) and following the manufacturer’s instructions with one minor modification: instead of the recommended 100 µL elution buffer, 70 µL of buffer was used for the elution to achieve a higher DNA concentration. DNA was extracted from 10 g aliquots of floor scrapes using the DNeasy PowerMax Soil Kit (10) (Cat# 12988-10) following the manufacturer’s instructions. The kit was chosen because of its ability to extract DNA from a large mass of soil and its design specifically tailored to soil samples with potentially low nucleic yield. DNA quantification was performed using a Qubit 4.0 Fluorometer. The initial volume of DNA elution was 5 mL, and initially, some samples yielded minimal DNA. However, by concentrating the volume to 200 µL using a 5M NaCl solution as suggested by the manufacturer, the DNA concentration increased significantly (Table A3), enabling successful sequencing for shotgun metagenomics. The extracted DNA quality was evaluated for suitability for subsequent shotgun sequencing using both Nanodrop and Qubit measurements. A Nanodrop spectrophotometer was used to assess DNA purity, where a 260/280 ratio reading of 1.8 indicated high quality and minimal contamination. Moreover, the 260/230 ratio reading was measured at 2.0, confirming minimal interference from substances such as carbohydrates or organic compounds.

### Library preparation and sequencing

300 ng DNA was used to prepare libraries by utilizing the Illumina DNA Prep Reagent Kit (Cat#20018705) and an automated liquid handler (epMotion 5075) according to the Illumina DNA Prep Reference Guide (1000000025416). The prepared DNA libraries underwent paired-end (2 x 150 bp) sequencing using the Illumina NextSeq550 platform (Illumina, San Diego, CA, USA).

### Taxonomical classification

We conducted pre-processing and taxonomical classification was completed using a previously published workflow.^84^ We performed multiple quality filtration steps to prepare sequencing reads for taxonomical analyses. We trimmed raw reads using TrimGalore v0.6.7 with a minimum quality score of 30 and minimum read length of 60, and deduplicated the trimmed reads using htstream SuperDeduper v1.3.0. For cow dung samples, we included an additional filter to exclude any reads that matched the cow host using a reference *Bos taurus (*cow) genome (ARS-UCD1.2) aligned with BWA v 0.7.17. We removed any unpaired (“orphan”) reads.

We completed taxonomic classification of the filtered reads using Kraken v2.1.3^85^ with the “PlusPF” index, which combines various reference sequences from the National Center for Biotechnology Information (NCBI) Reference Sequence Database (RefSeq) that include archaea, bacteria, viral, plasmid, human, protozoa and fungi genomes. Kraken results were passed through Bracken v2.9.0 to estimate abundance at every taxonomic level. In some analyses, we restricted Bracken outputs to organisms with known pathogenicity in humans^38^.

### Antibiotic resistance genes

We used the CZID platform to identify ARGs in both sample types. We downloaded report tables from the AMR module and filtered to ARGs with read coverage breadth > 10% or contig coverage breadth > 10% and > 5 reads mapped^86^. Because short reads may not adequately differentiate between highly homologous alleles or genes belonging to the same gene family, we restricted to one gene per sample^86^. For AMR detection, the Resistance Gene Identifier (RGI) tool was used to compare filtered reads and contigs against AMR reference sequences from the Comprehensive Antibiotic Resistance Database (CARD)^87,88^. When ARGs were identified using both the read and contig methods within a single sample, only the ARG identified using contigs was included. We generated a heatmap of ARGs within each sample and classified each ARG by a categorical variable indicating contigs match quality (perfect, strict, nudged) or read matches. “Perfect” contig matches identically matched reference sequences in CARD^89^. “Strict” contig matches were those that matched previously unknown variants of known ARGs, including secondary screening for key mutations. “Nudged” contig matches had at least 95% identity to known AMR genes and were matched using a percent identity threshold not taking alignment length into account. Heatmaps show the highest confidence level for contig matches for each ARG in each sample.

### Statistical analysis

We determined the alpha-diversity within each sample type using the richness attribute, Chao index for species richness^39–41^, Pielou’s evenness index^42^, Shannon index^43^, Simpson index^44^, and inverse Simpson index. We tested for differences in alpha-diversity between sample types using the Wilcoxon signed-rank test^90^. To assess microbial diversity, we calculated the Bray Curtis dissimilarity^91^ and investigated clustering by sample type, household membership, and the presence of visible animal feces on floors at the time of measurement using principal coordinates analysis^92^ and permutational multivariate analysis of variance (PERMANOVA)^93^. Alpha and beta diversity metrics were calculated using the R package *vegan*^94^.

### Ethics

The parent study was approved by the International Centre for Diarrhoeal Disease Research, Bangladesh Ethical Review Committee (PR-22069) and the Stanford Institutional Review Board (63990). Participants provided written informed consent.

### Data Availability

Raw sequencing data are deposited under the NCBI Bioproject PRJNA1130536. URL for reviewers: https://dataview.ncbi.nlm.nih.gov/object/PRJNA1130536?reviewer=leg1d9ic67vo6ugmc7jkf407li Analysis scripts and files are available at https://github.com/kalanir/cradle-pilot-seq.

## Appendix Tables and Figures

**Table A1.**
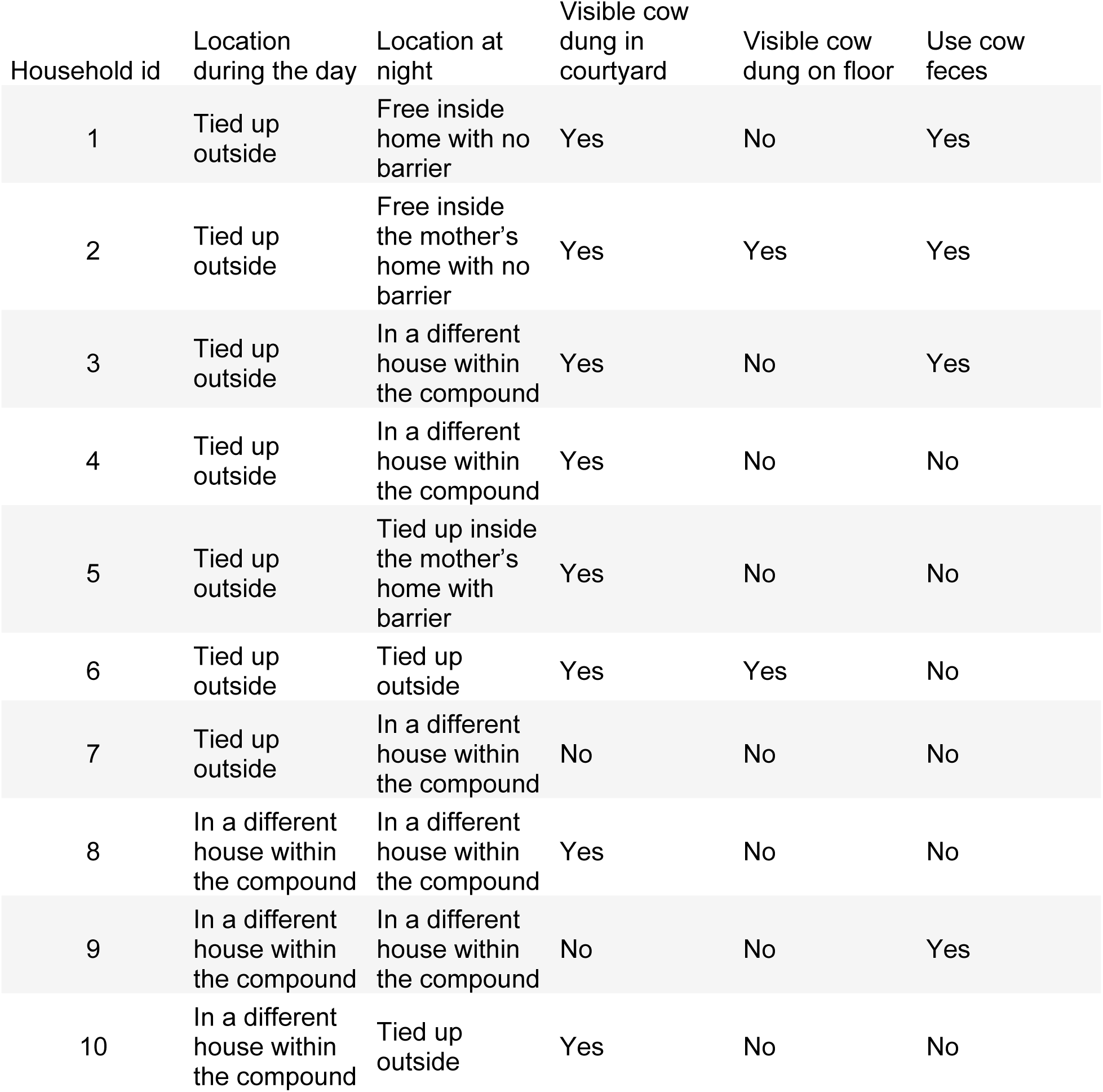
Cow-related characteristics of each household

| Household id | Location during the day | Location at night | Visible cow dung in courtyard | Visible cow dung on floor | Use cow feces |
| --- | --- | --- | --- | --- | --- |
| 1 | Tied up outside | Free inside home with no barrier | Yes | No | Yes |
| 2 | Tied up outside | Free inside the mother's home with no barrier | Yes | Yes | Yes |
| 3 | Tied up outside | In a different house within the compound | Yes | No | Yes |
| 4 | Tied up outside | In a different house within the compound | Yes | No | No |
| 5 | Tied up outside | Tied up inside the mother's home with barrier | Yes | No | No |
| 6 | Tied up outside | Tied up outside | Yes | Yes | No |
| 7 | Tied up outside | In a different house within the compound | No | No | No |
| 8 | In a different house within the compound | In a different house within the compound | Yes | No | No |
| 9 | In a different house within the compound | In a different house within the compound | No | No | Yes |
| 10 | In a different house within the compound | Tied up outside | Yes | No | No |

**Table A2.**
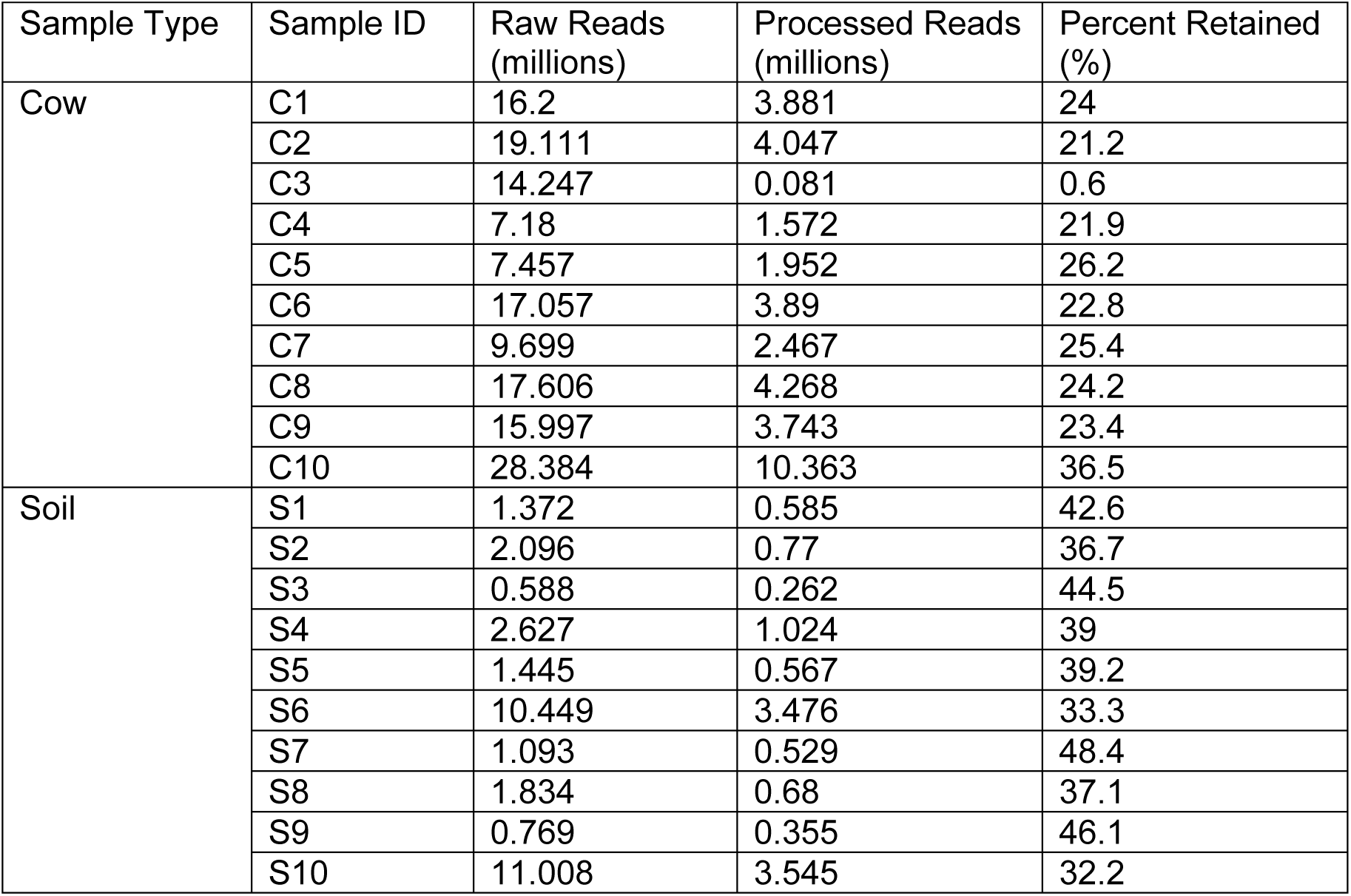
Raw and processed read counts for each sample

| Sample Type | Sample ID | Raw Reads<br>(millions) | Processed Reads<br>(millions) | Percent Retained<br>(%) |
| --- | --- | --- | --- | --- |
| Cow | C1 | 16.2 | 3.881 | 24 |
|  | C2 | 19.111 | 4.047 | 21.2 |
|  | C3 | 14.247 | 0.081 | 0.6 |
|  | C4 | 7.18 | 1.572 | 21.9 |
|  | C5 | 7.457 | 1.952 | 26.2 |
|  | C6 | 17.057 | 3.89 | 22.8 |
|  | C7 | 9.699 | 2.467 | 25.4 |
|  | C8 | 17.606 | 4.268 | 24.2 |
|  | C9 | 15.997 | 3.743 | 23.4 |
|  | C10 | 28.384 | 10.363 | 36.5 |
| Soil | S1 | 1.372 | 0.585 | 42.6 |
|  | S2 | 2.096 | 0.77 | 36.7 |
|  | S3 | 0.588 | 0.262 | 44.5 |
|  | S4 | 2.627 | 1.024 | 39 |
|  | S5 | 1.445 | 0.567 | 39.2 |
|  | S6 | 10.449 | 3.476 | 33.3 |
|  | S7 | 1.093 | 0.529 | 48.4 |
|  | S8 | 1.834 | 0.68 | 37.1 |
|  | S9 | 0.769 | 0.355 | 46.1 |
|  | S10 | 11.008 | 3.545 | 32.2 |

**Table A3.** Nucleic acid yield for each sample

| Household<br>id | Soil floor sample<br>ng/μL | Cow dung sample<br>ng/μL |
| --- | --- | --- |
| 1 | 34.9 | 237 |
| 2 | 36.5 | 186 |
| 3 | 32.8 | 228 |
| 4 | 36.5 | 208 |
| 5 | 43.1 | 298 |
| 6 | 38.9 | 320 |
| 7 | 29.7 | 212 |
| 8 | 39.3 | 237 |
| 9 | 37.3 | 197 |
| 10 | 35.2 | 282 |

**Figure A1.**
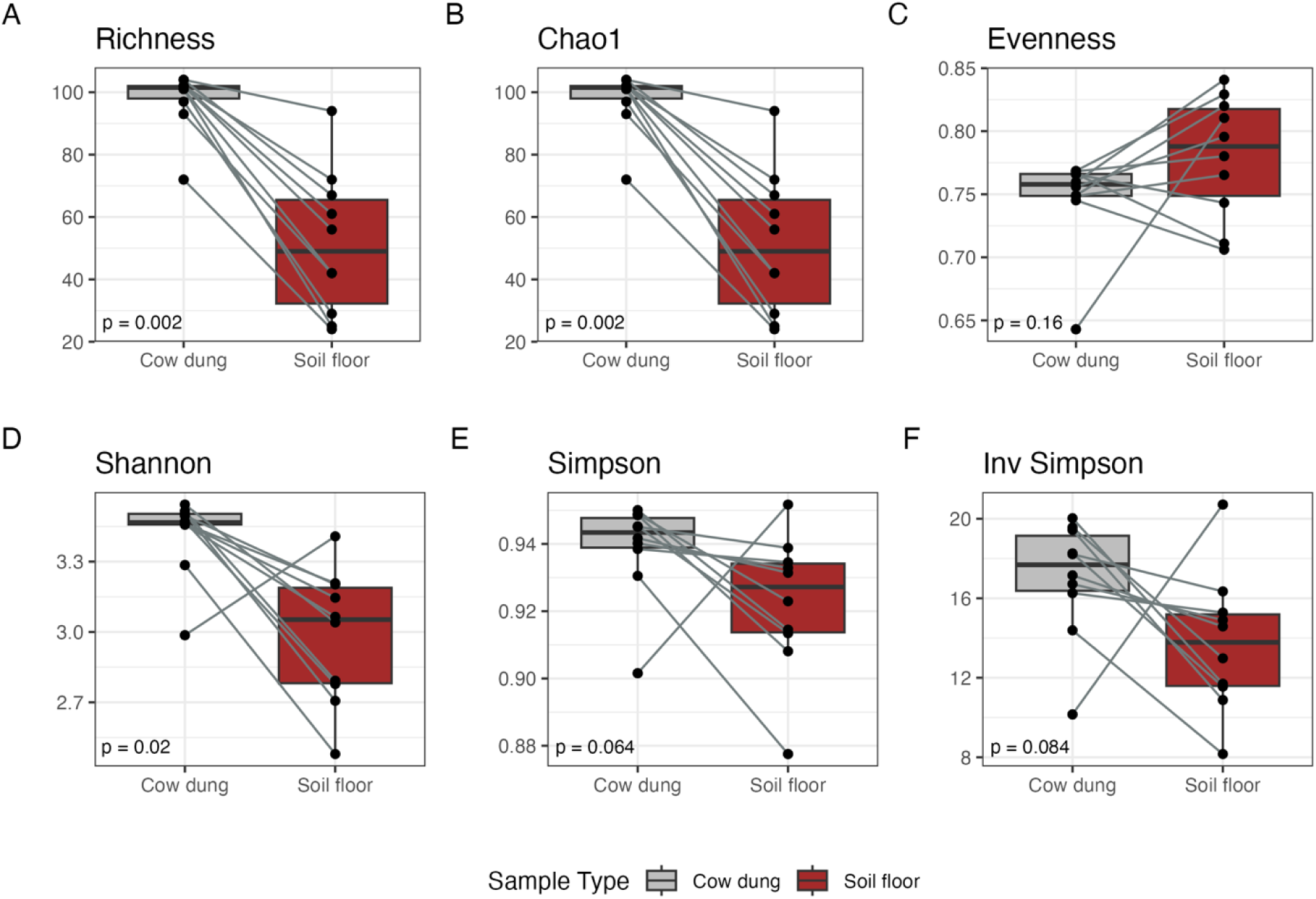
Alpha-diversity indices for potential pathogen genera by sample type. Includes 10 paired cow dung and soil floor samples. Indices were compared between sample types using the Wilcoxon signed-rank test.

**Figure A2.**
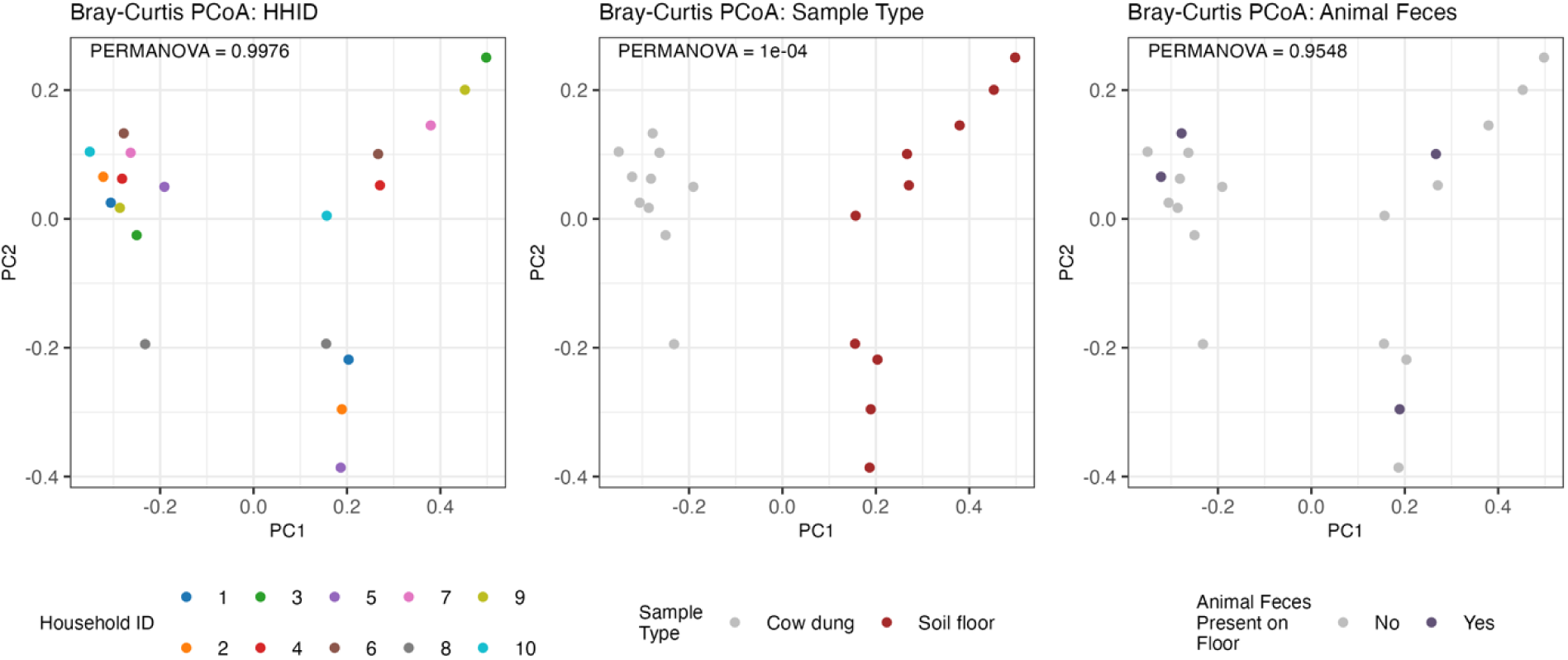
Bray Curtis dissimilarity between communities of potential pathogen species by household membership, sample type, and presence of animal feces on the household floor. Includes 10 household-paired cow dung and soil floor samples. Bray-Curtis dissimilarity was compared between sample types using PERMANOVA.

**Figure A3.**
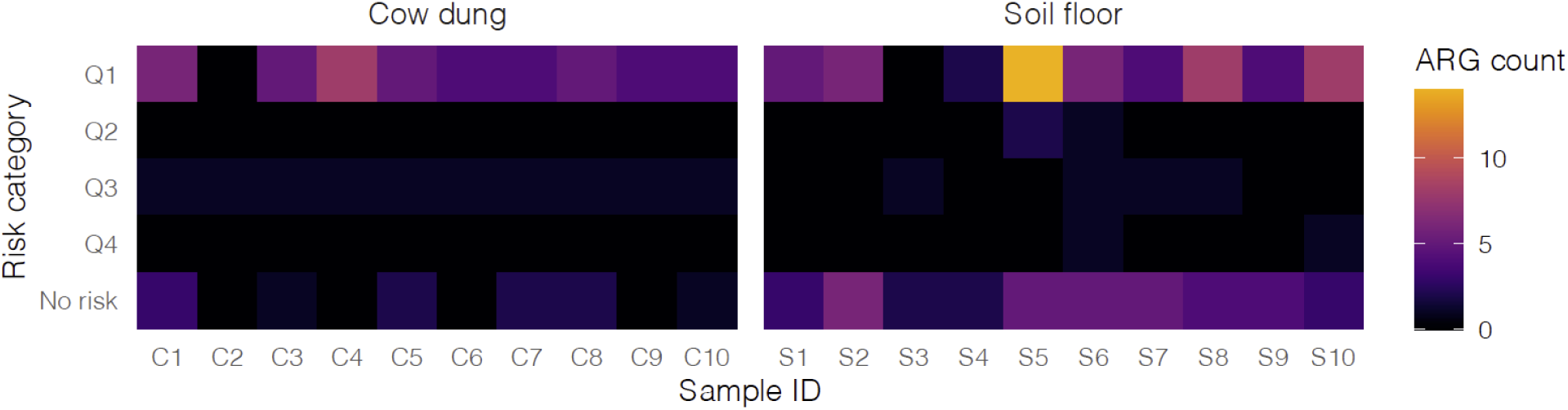
Number of antibiotic resistance genes (ARGs) in each human health risk quartile in each sample. Based on classifications in Zhang et al., 2022 (https://doi.org/10.1038/s41467-022-29283-8).

## Acknowledgements

The funders had no role in study design, data collection and interpretation, or the decision to submit the work for publication. This study was funded by a grant from the National Institute of Child Health and Human Development to Stanford University (R01HD108196; PI: Benjamin-Chung) and National Institute of Allergy and Infectious Diseases (F31AI179107; PI: Nguyen). ATN was also supported by the Stanford Data Science Scholars Program. JBC is a Chan Zuckerberg Biohub Investigator. The content is solely the responsibility of the authors and does not necessarily represent the official views of the National Institutes of Health. Some of the computing for this project was performed on the Sherlock cluster. We would like to thank Stanford University and the Stanford Research Computing Center for providing computational resources and support that contributed to these research results.

Author contributions: Conceptualization (JBC, AE, IS, FJ, MR), Data curation (AKS, GBH), Formal analysis (KR, ATN, JBC), Funding acquisition (JBC, MR, AE), Investigation (Mustafizur R, MJ, ST, AY, SH, ZHM, AKS, FJ), Methodology (JBC, AE, KR, ATN, CA, MMR, MJ), Project administration (SH, AY, FJ), Resources (Mustafizur R, MJ), Software (JBC, KR, GBH), Supervision (JBC, AE, FJ, MR, Mustafizur R), Validation (CA, GBH), Visualization (KR, ATN, GBH, CA, JBC), Writing - original draft (ATN, KR, JBC), Writing - review & editing (all authors)

